# The evolution, distribution and diversity and of endogenous circoviral elements

**DOI:** 10.1101/207399

**Authors:** Tristan P.W Dennis, William Marciel de Souza, Soledad Marsile-Medun, Joshua B. Singer, Sam J. Wilson, Robert J. Gifford

## Abstract

Circoviruses (family *Circoviridae*) are small, non-enveloped viruses that have short, single-stranded DNA genomes. Circovirus sequences are frequently recovered in metagenomic investigations, indicating that these viruses are widespread, yet they remain relatively poorly understood. Endogenous circoviral elements (CVe) are DNA sequences derived from circoviruses that occur in vertebrate genomes. CVe can provide unique, retrospective information about the biology and evolution of circoviruses. In this study, we screened 362 vertebrate genome assemblies *in silico* to generate a catalog of CVe loci. We identified a total of 179 CVe sequences, most of which have not been reported previously. We show that these CVe loci reflect at least 19 distinct germline integration events. We determine the structure of CVe loci, identifying some that show evidence of potential functionalization. We also identify orthologous copies of CVe in snakes, fish, birds, and mammals, allowing us to add new calibrations to the timeline of circovirus evolution. Finally, we observed that some ancient CVe group robustly with contemporary circoviruses in phylogenies, with all sequences within these groups being derived from the same host class or order, implying a hitherto underappreciated stability in circovirus-host relationships. The openly available dataset constructed in this investigation provides new insights into circovirus evolution, and can be used to facilitate further studies of circoviruses and CVe.

**Abbreviations:**

CVe: endogenous circoviral element
ORF: open reading frame
Cap: Capsid
Rep: Replicase
ssDNA: single-stranded DNA

## 1. Introduction

Circoviruses (family *Circoviridae*, genus Circovirus) are small, non-enveloped viruses with single-stranded DNA (ssDNA) genomes. Circovirus genomes are typically ~2 kilobases (kb) in length and contain only two open reading frames (ORFs): one encoding a non-structural replicase (Rep) protein, and a second encoding the viral capsid (Cap). The family contains two genera: *Circovirus* and *Cyclovirus* [1]. Over recent years, sequencing-based virus discovery efforts have identified many new members of these two genera [2]. However, little or nothing is known about most of the novel viruses that have been identified using these approaches. Only a handful of circoviruses have been investigated at a level beyond sequencing: porcine circoviruses 1 and 2 (PCV-1 and PCV-2), which infect swine, and beak and feather disease virus (BFDV), which infects various avian species [3].

Circovirus sequences are frequently recovered from tissue and environmental samples in metagenomic investigations, indicating that these viruses are widespread, yet they remain relatively poorly understood [4]. Endogenous circoviruses (CVe) provide an unconventional but useful source of information about circovirus distribution, diversity and evolution. These sequences are derived from the genomes of circoviruses that circulated millions of years ago, and became integrated into the host germline [5, 6]. Relatively robust minimum age estimates can be obtained for CVe via the identification of orthologous copies in distinct host lineages. On this basis, we now know that the association between circoviruses and vertebrates extends back millions of years before the present day [7, 8].

In this study, we screened vertebrate genomes *in silico* to generate a comprehensive catalog of CVe. We used these data to: (i) extract information about the long-term evolution of circoviruses; (ii) generate an openly accessible data resource that can facilitate the further investigation of CVe and circoviruses.

## 2. Material & Methods

### 2.1. Identification and analysis of CVe sequences

We used similarity searches to systematically screen genome assemblies of 362 chordate species (**Table S1**) for sequences homologous to circovirus proteins. Vertebrate genome assemblies and circovirus reference genomes were obtained from the NCBI genomes resource. Screening *in silico* was performed using the database-integrated genome-screening tool. The DIGS procedure used to identify CVe comprises two steps. In the first, a circovirus probe sequence (e.g. a Cap or Rep protein sequence) is used to search a particular genome assembly file using the basic local alignment search tool (BLAST) program [9]. In the second, sequences that produce statistically significant matches to the probe are extracted and classified by BLAST-based comparison to a set of virus reference genomes (see **Table S2**). Results are captured in a MySQL database.

We inferred the ancestral ORFs of CVe (and the number of stop codons and frameshifts interrupting these ORFs) via a combination of automated alignment and manual adjustment. Multiple sequence alignments were constructed using MUSCLE [10] and PAL2NAL [11]. Manual inspection and adjustment of alignments was performed in Se-Al [12]. Phylogenies were constructed using maximum likelihood as implemented in RaxML [13], and the VT protein substitution model [14] as selected using ProTest [15].

### 2.2. Construction of CVe sequence data resource

We used GLUE - an open, data-centric software environment specialized in capturing and processing virus genome sequence datasets – to collate the sequences, alignments and associated data used in this investigation. The aim was to create a standardized data CVe resource that would be openly accessible, and would facilitate the further use and development of the dataset assembled here. The project includes all the CVe identified by our *in silico* screen, as well as a set of representative reference sequences for the *Circovirus* genus (**Table S2**). All of these sequences are linked to the appropriate auxiliary data; for the virus sequences, this includes information about the sample from which the sequence was obtained; for CVe, it includes the name of genome assembly and contig in which the CVe sequence was identified, and its coordinates and orientation within that contig.

The project also includes the key alignments constructed in this study, linked together using the GLUE ‘alignment tree’ data structure. These include: (i) ‘tip’ alignments in which all taxa are CVe that are known or putative orthologs of one another; (ii) a ‘root’ alignment constructed to represent proposed homologies between the genomes of representative viruses in the genus *Circovirus* and the CVe recovered by our screen. Because each of these alignments is constrained to a standard reference sequence, are alignments are linked to one another.

We applied a systematic approach to naming CVe. Each element was assigned a unique identifier (ID) constructed from a defined set of components. The first component is the classifier ‘CVe’. The second is a composite of two distinct subcomponents separated by a period: the name of CVe group (usually derived from the host group in which the element occurs in (e.g. Carnivora), and the second is a numeric ID that uniquely identifies the insertion. Orthologous copies in different species are given the same number, but are differentiated using the third component of the ID that uniquely identifies the species from which the sequence was obtained. An additional unique numeric ID may be added to this component in cases were a CVe element has expanded via duplication.

## 3. Results

### 3.1. Identification and phylogenetic analysis of vertebrate CVe

We systematically screened 362 vertebrate genome assemblies for CVe, and identified a total of 179 CVe sequences (**Table S3**), in 52 distinct species (**Table 2**). For each CVe sequence, we determined the regions of the circovirus genome represented, and attempted to identify genomic flanks. Where genomic flanks were present, we compared these with one another to identify potentially orthologous CVe loci. In several cases, it was not possible to determine whether multiple CVe loci within the same species (or group of closely-related species) represented the outcome of distinct incorporation events, or the fragmented remains of a single, ancestrally acquired element. The main causes of uncertainty were; (i) lack of flanking sequences due to short contig length, or undetermined DNA sequences flanking CVe, and; (ii) the presence of multiple CVe that spanned non-overlapping regions of the circovirus genome. Since CVe are comparatively rare in vertebrate genomes [7, 16], we conservatively assumed a single incorporation event had taken place except in cases where it could be demonstrated otherwise. On this basis, we estimate that the 179 CVe identified here represent at least 19-26 distinct germline incorporation events (**Table 1**, **Table 2**, **Figure 1**), depending on whether CVe in ray-finned fish are taken to represent a single incorporation event, or seven distinct incorporation events, each in a different order (see **section 3.3**). The large discrepancy between the number of elements versus the number of events reflects the fact that 101 of the 179 CVe identified in our study (57%) belong to a group of highly duplicated CVe loci in carnivore genomes, all of which derive from a single germline incorporation event.

**Table 1.** CVe detected in published vertebrate genome assemblies.

| EVE name | Reference | Genes | # Seqs | # Species |
| --- | --- | --- | --- | --- |
| **Agnatha** |  |  |  |  |
| CVe- *Eptatretus* * | This study | Rep | 7 | 1 |
| **Bony Fish** |  |  |  |  |
| CVe- *Anguilla* * | This study | Rep |  | 1 |
| CVe- *Characiformes* * | This study | Rep |  | 1 |
| CVe- *Clupeiformes* * | This study | Rep |  | 1 |
| CVe-*Cypriniformes* | [19] | Rep-Cap |  | 2 |
| CVe- *Cyprinodontiformes* * | This study | Rep |  | 1 |
| CVe- *Perciformes* * | This study | Rep |  | 3 |
| CVe- *Salmoniformes* * | This study | Rep |  | 1 |
| **Amphibians** |  |  |  |  |
| CVe-*Anura* | [16] | Rep | 2 | 2 |
| **Reptiles** |  |  |  |  |
| CVe-*Viperidae* | [23] | Rep-Cap | 16 | 6 |
| **Birds** |  |  |  |  |
| CVe-*Tinamou* | [16] | Rep-Cap | 2 | 1 |
| CVe-*Psittaciformes* | [16] | Rep-Cap | 4 | 3 |
| CVe-*Passeriformes* | [16] | Rep | 7 | 5 |
| CVe-*Egretta* | [16] | Rep | 1 | 1 |
| CVe- *Gallirallus* * | This study | Rep | 1 | 1 |
| CVe- *Picoides* * | This study | Rep | 1 | 1 |
| **Mammals** |  |  |  |  |
| CVe- *Chrysochloris* * | This study | Rep, Cap | 3 | 1 |
| CVe-*Carnivora* | [7] | Rep | 101 | 13 |
| CVe- *Mus.caroli* * | This study | Rep | 1 | 1 |
| CVe-*Heterocephalus* * | This study | Rep | 1 | 1 |
| CVe- *Phascolarctos* * | This study | Rep | 1 | 1 |
| CVe-*Sarcophilus* * | This study | Rep | 1 | 1 |
| CVe-*Monodelphis* |  | Rep | 1 | 1 |
| CVe-*Galeopterus* * | This study | Rep | 2 | 1 |
| CVe- *Manis* * | This study | Rep | 1 | 1 |
| CVe- *Choloepus* * | This study | Cap | 2 | 1 |
| Totals |  |  | 179 | 53 |

3 **Footnote:** Asterisks indicate newly identified CVe loci.

**Table 2.**
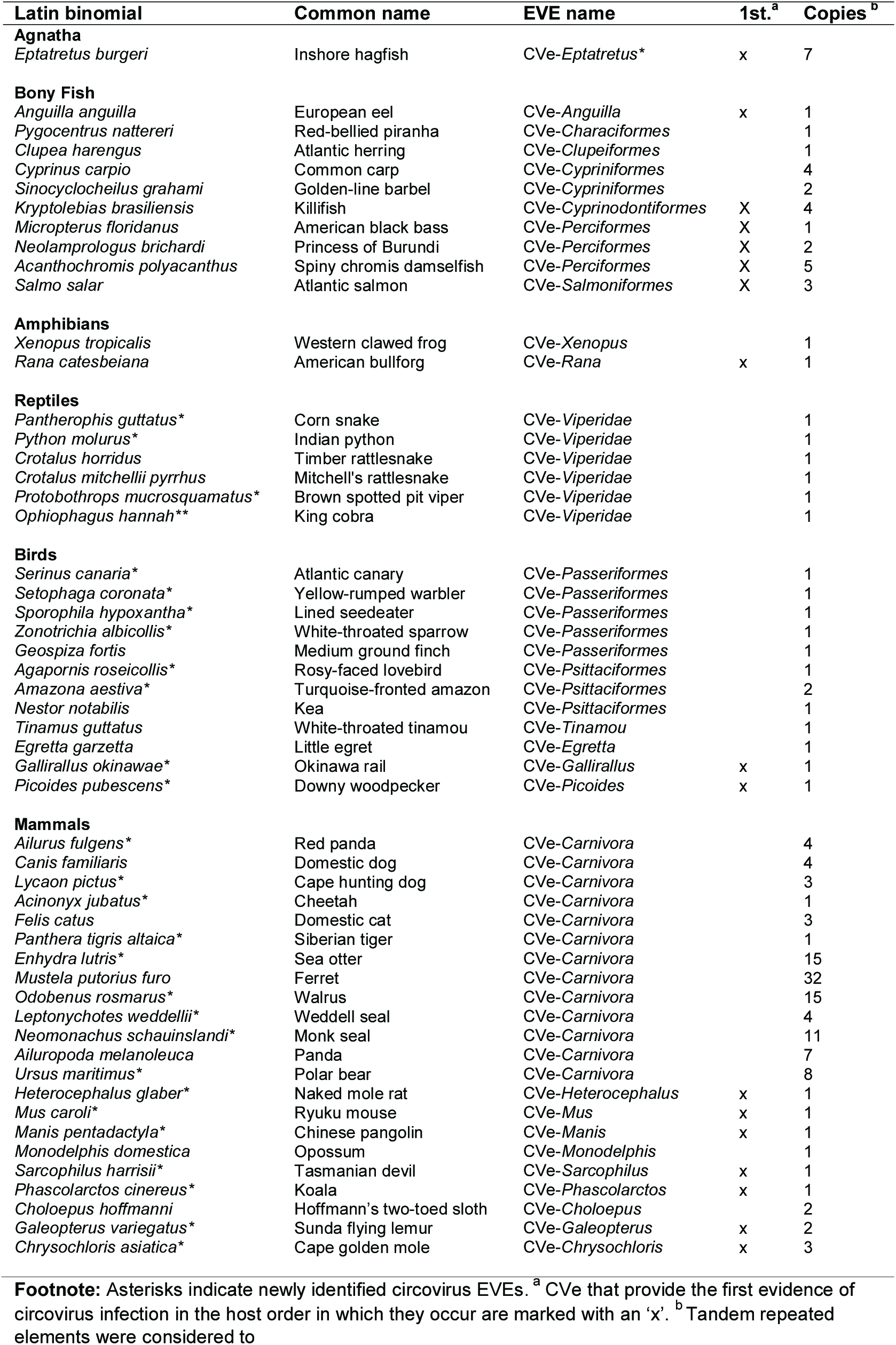
Vertebrate species with CVe.

**Figure 1.**
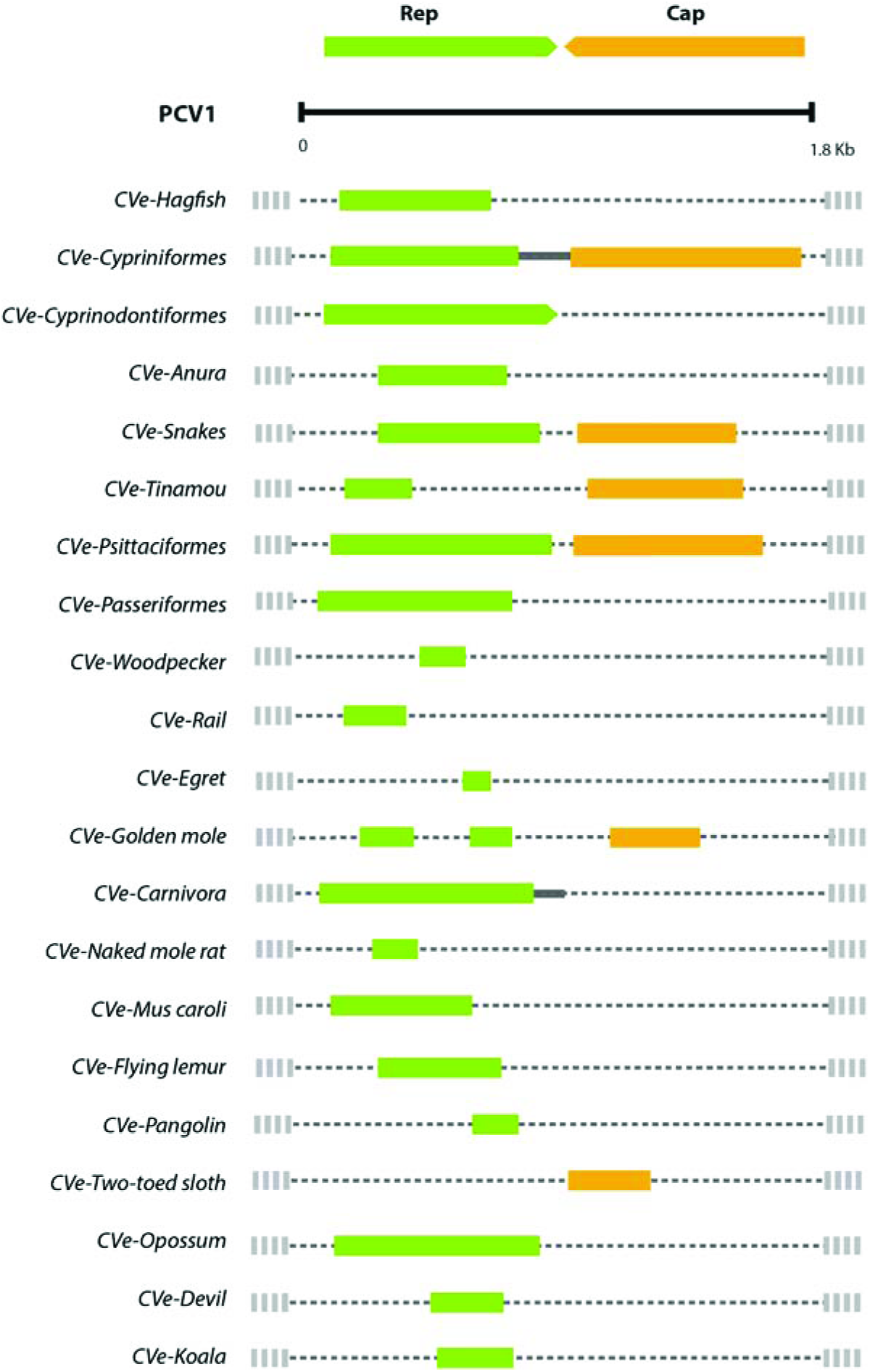
Genome structures of 21 endogenous circovirus (CVe) elements identified in vertebrate genomes. CVe coding sequences are represented schematically as green and yellow bars relative to a porcine circovirus 1 (PCV1) reference genome (accession # NC_001792.2).

We only identified four cases where CVe encoding both *rep* and *cap* were present in the same species or species group. In most, only *rep*-derived sequences appear to have been incorporated/retained, and in one case only *cap* (**Table 1**). We constructed a multiple sequence alignment (MSA) that spanned the entire circovirus genome and contained both reference sequences for CVe (these could be based on individual loci, or a consensus), and representative circovirus reference taxa (**Table S2**). We used this ‘root’ MSA (see **section 2.2**) to infer which regions of the circovirus genome had been incorporated as CVe. Where CVe spanned coding sequence, we inferred the putative ancestral reading frame by comparing CVe and circovirus sequences, and attempting to identify likely frameshifting mutations. Most CVe represent only fragments of the genome (**Figure 1**), and many are relatively degraded, containing multiple frameshifting indels and stop codons.

Where we identified several CVe from the same species, we compared genomic regions to search for evidence of homology and thereby identify orthologs. Where we were able to identify orthologous CVe insertions, we used these data to create a timeline of circovirus evolution (**Figure 2**). In addition, we identified sets of ‘potentially orthologous’ CVe, where sequence similarity and phylogenetic relationships were consistent with orthology, but this could not be confirmed or ruled out based on flanking sequences.

**Figure 2.**
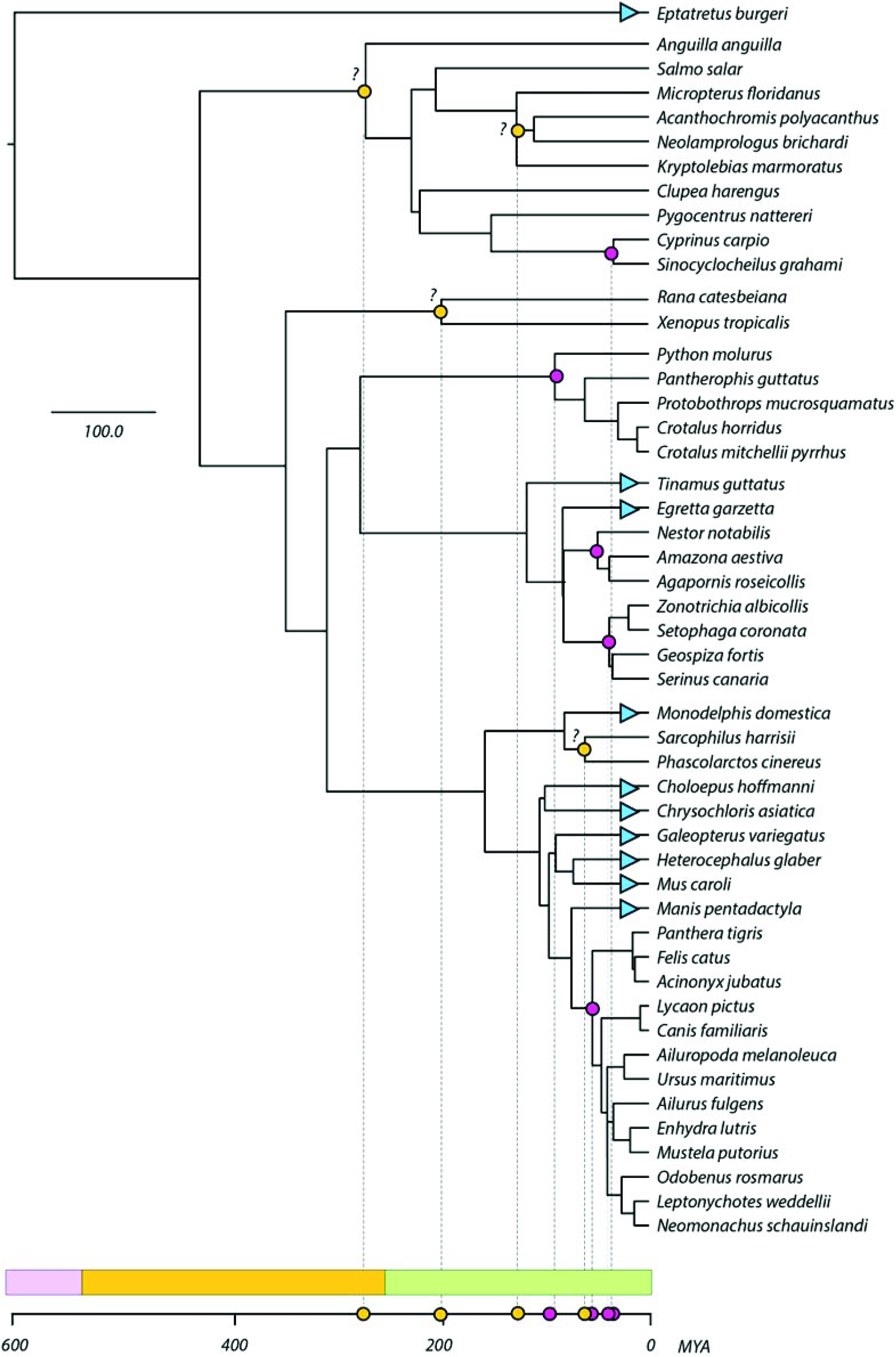
Evolutionary relationships of vertebrate species in which CVe have been identified, and timeline of CVe evolution. Pink circles indicate confirmed orthologs. Yellow circles indicate the presence of potential orthologs that have not been confirmed. Blue triangl2es indicate where CVe loci are present, but no information about their ages could be obtained.

A range of distinct partitions were derived from the virtually translated root MSA (with frameshifts removed), and used to construct bootstrapped ML phylogenies (**Figure 3**). In general, support for the deeper branching relationships between CVe and circoviruses was weak, irrespective of which genomic region was used to construct trees. This reflects the fact that most CVe are short and/or highly degraded, and these sequences tend to group distantly from other taxa. However, in phylogenies based on Rep (**Figure 3**), several robustly supported subgroupings were observed, three of which – referred to here as mammal 1, cyprinid 1, and avian 1-included a mixture of CVe and contemporary circoviruses. Notably, in all three of these clades, CVe and circovirus sequences were obtained from the same hosts of the same taxonomic class. The sections that follow describe the distribution and diversity of CVe within individual classes in the subphylum Vertebrata (chordates with backbones).

**Figure 3.**
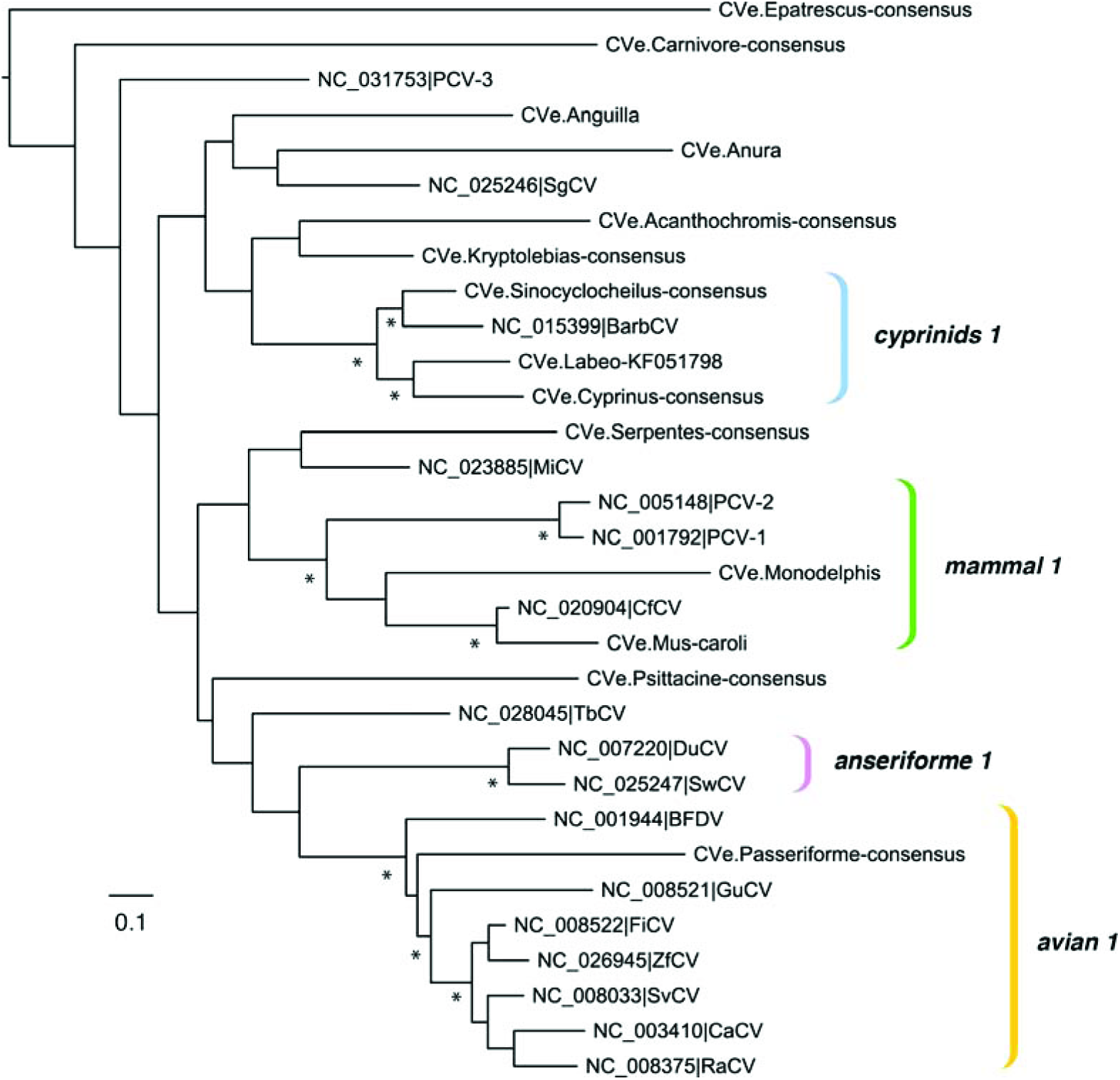
A maximum likelihood phylogeny showing estimated evolutionary relationships between endogenous circovirus (CVe) elements and exogenous circoviruses. The phylogeny constructed from an alignment spanning ~200 amino acids in Rep. The scale bar shows evolutionary distance in substitutions per site.

### 3.2. CVe in jawless vertebrates

Extant vertebrates are divided into the jawed vertebrates (Gnathostomata) and jawless vertebrates (Agnatha). The Agnatha represent the most basal group of vertebrates and includes the hagfishes (myxinoids) and lampreys (petromyzontids). We identified seven sequences exhibiting homology to *rep* in the genome assembly of the inshore hagfish (*Eptatretus burgeri*). These sequences are relatively distinct from other circoviruses, and also showed relatively high genetic diversity relative to one another, forming three distinct groups in phylogenetic trees (**Figure S1**). Notably, the putative Rep polypeptides encoded by these sequences contained several in-frame indels relative to one another. Because such a pattern of variation is unlikely to arise through neutral accumulation of mutations in the germline, this suggests the occurrence of at least three distinct genome incorporation events, each involving distinct, but relatively closely related viruses. However, since we were unable to identify unambiguous genomic flanking sequences for any of these loci, their classification as CVe should for now be considered tentative.

### 3.3. CVe in ray-finned fish (class Actinopterygii)

Circoviruses are thought to infect barbel fish (*Barbus barbus*) and European catfish (*Silurus glanis*), based on (i) the observation of viral particles in tissues, and the recovery of circovirus sequences from these tissues via nested PCR [17, 18]. In addition, CVe have been reported in one fish species - the Indian rohu (*Labeo rohita*) [19]. We identified numerous additional CVe sequence in the genome assemblies of ray-finned fishes (Class *Actinopterygii*) (**Table 2, Table S3**). We established that at least two of these CVe - occurring in the common carp (*Cyprinus carpio*) and golden-line barbell (*Sinocyclocheilus grahami*) genomes - were orthologs of one another, indicating they were incorporated into the germline of cyprinid fish more than 39 million years ago [28, 29]. These CVe were comprised of multiple complete circovirus genomes arranged in tandem, and intriguingly, were observed CVe group as sister taxa to barbel circovirus (BarbCV) in phylogenetic trees, sharing ~70% nucleotide identity (across 1654 nucleotides) with the BarbCV genome.

We also identified matches to *rep* in eight other species of ray-finned fish (**Table 2**). We could not determine with certainty how many integration events these CVe represented. Interestingly, however, all of these sequences group together in phylogenies (**Figure 3**), and the phylogeny constructed for these elements - when rooted on the CVe from the most basal host - the European eel (*Anguilla anguilla*), approximately follows that of the host species, consistent with a single ancestral integration event >200 million years ago (**Figure 2**). Alternatively, the CVe observed in distinct orders might represent distinct incorporation events. This is supported by the placement of CVe.anura in phylogenies, in which it splits the fish CVe from one another, albeit with weak support (**Figure S1**). In addition, the observation that CVe elements in order cypriniforme fish (golden-line barbell and carp) occur as full-length tandem genomes, whereas those in Perciformes are derived from more divergent fragments of *rep*, is suggestive of at least two separate incorporation events. Notably one CVe in the mangrove rivulus (*Kryptolebias marmoratus*) encoded a complete intact *rep* gene (**Figure 1**) that is predicted to be expressed, suggesting it may have been functionalized in some manner.

### 3.4. CVe in amphibians

Sequences homologous to circovirus *rep* genes have previously been identified in the Western clawed frog (*Xenopus tropicalis*) [20]. We identified CVe in the genome of the American bullfrog (*Rana catesbeiana*) that partially overlaps that identified in *Xenopus*. Potentially, these sequences could be orthologs of one another, which would imply a minimum age of ~204 MYA [21, 22] (**Figure 2**). However, we were unable to confirm this based on analysis of flanking genomic sequences.

### 3.5. CVe in reptiles

A pair of othologous CVe, each covering about 75% of the circovirus genome, have previously been recovered from rattlesnake genomes (*Crotalus spp*) [23]. We identified CVe in four additional snake species (**Table 2**). Examination of aligned snake CVe sequences indicated that all are likely to be orthologs of those previously reported in rattlesnakes (see **Figure S2**), implying that this CVe integrated into the serpentine germline ~72-90 million years ago (Mya) (**Figure 2**).

### 3.6. CVe in birds (class Aves)

CVe have previously been reported in the genomes of several avian species: the little egret (*Egretta garzetta*), white-throated tinamou (*Tinamus guttatus*), medium ground-finch (*Geospiza fortis*), and kea (*Nestor notabilis*) [16, 20]. We identified CVe in eight additional species. Some of these appeared likely to be orthologs of CVe reported previously. For example, we identified CVe in two species of psittacine bird that appeared represented orthologs of one another, and possibly of those previously identified in the kea (*Nestor notabilis*) [16] (**Table 2**), which would imply integration into the psittacine germline prior to the divergence of the major extant lineages within the order Psittaciformes (estimated to have occurred 30-60 Mya [13, 24]) (**Figure 2**).

We also identified orthologs of the *rep*-derived insertion previously described in the medium ground finch in several additional species in the avian order Passeriformes (songbirds) (**Table 2**). Identification of these orthologs demonstrates that this particular CVe predates the radiation of avian sub-order Passeroida ~38 Mya [13, 25] (**Figure 2**).

In addition to identifying the previously reported CVe in the genomes of the white-throated tinamou (*Tinamus guttatus*) and little egret (*Egretta garzetta*) [16], we identified previously unreported CVe in the Japanese rail (*Gallirallus okinawae*: order Gruiformes) and downy woodpecker (*Picoides pubescens*: order Piciformes) (**Table 2**). Both these sequences were relatively short and divergent, and consequently we could not determine their relationships to other CVe and circoviruses with confidence.

### 3.6. CVe in mammals (class Mammalia)

The majority of CVe identified in our screen were recovered from carnivore genome assemblies. As far as we are able to discern from phylogenetic and comparative analysis, all of these CVe derive from a 1-4 germline incorporation events involving an ancient carnivore *rep* gene. However, the copy number of these elements has expanded subsequent to their incorporation into the germline, in some cases quite dramatically. The grouping of carnivore CVe in phylogenies (**Figure 4**) indicates that at least four CVe insertions were present in the carnivore germline prior to the divergence of extant families within this order. The copy number of one particular element (referred to here as CVe-Carnivora-4) has expanded in some carnivore lineages. As shown in **Figure 4**, the phylogenetic relationships between duplicates in the group CVe-Carnivora-4 indicate that these expansions have occurred independently in ursids (bears), pinnipeds (seals and walruses), and mustelids. CVe in this lineage are flanked by sequences that disclose homology to non-LTR retrotransposons. Thus, one plausible explanation for the elevated copy number in certain carnivore lineages is that CVe have become embedded into retroelements and copied along with these sequences when they undergo transposition.

**Figure 4.**
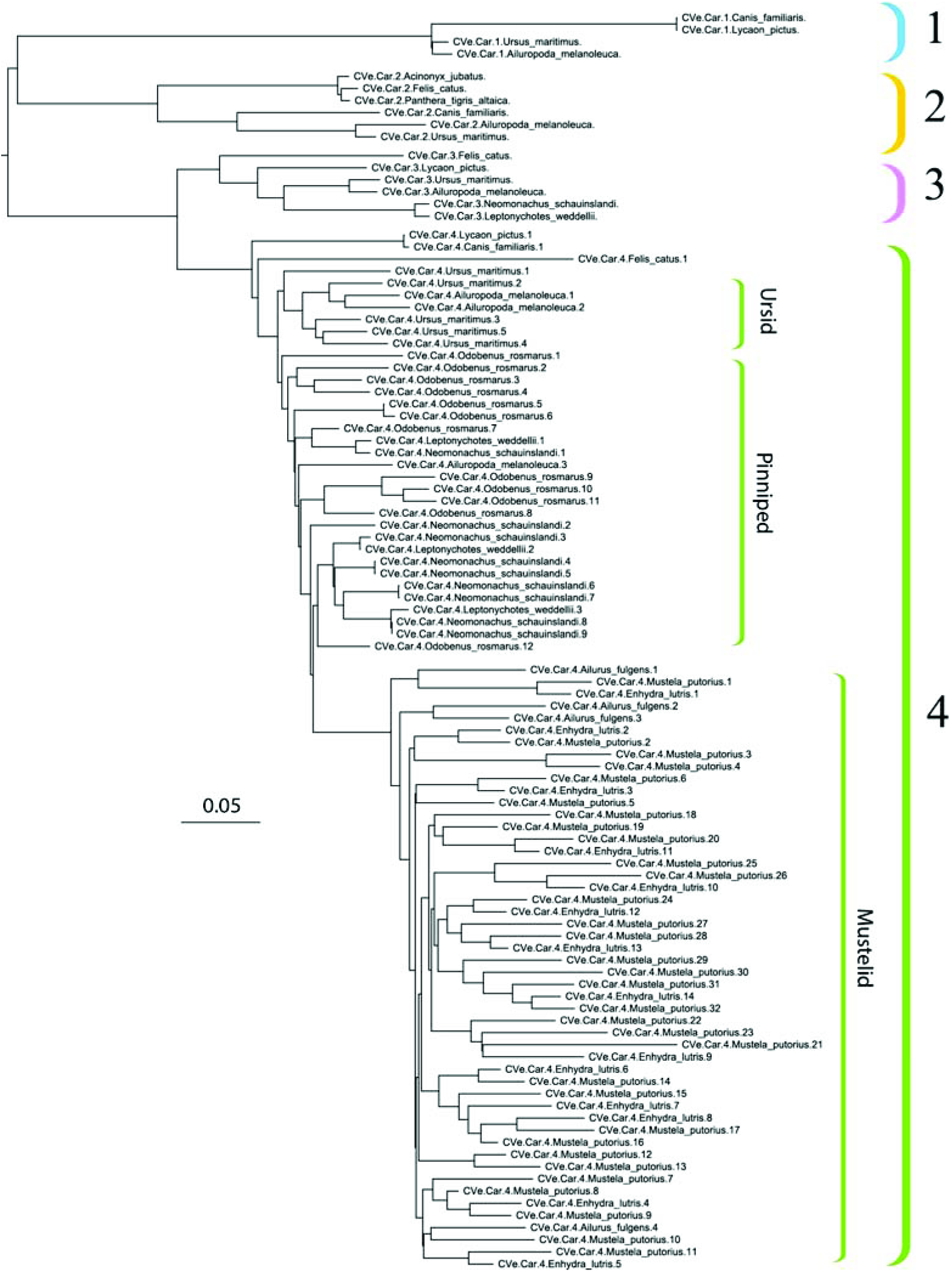
Phylogeny of CVe sequences recovered from carnivore genome assemblies. At least four distinct CVe loci are present in the carnivore germine (clades I-IV) as indicated by the coloured brackets. Within group IV, distinct copy number expansions appear to have occurred in ursids (bears), pinnipeds (seals and walruses), and mustelids. The scale bar shows evolutionary distance in substitutions per site. The tree is midpoint rooted for display purposes.

A novel, relatively well-preserved *rep*-derived CVe was identified in the genome of the Ryukyu mouse (*Mus caroli*) that grouped closely with circoviruses genome recovered from dogs [26, 27]. This element presumably arose after this species diverged from the house mouse (*Mus musculus*) ~6-7 Mya, since it is absent from this species.

In the cape golden mole (*Chrysochloris asiaticus*) matches to both *cap* and *rep* were identified. However, these occurred on distinct contigs and did not overlap. Furthermore, both CVe were relatively short and degraded, and were highly divergent relative to other CVe. CVe derived from *cap* were also identified in the genome of Hoffmann’s two-toed sloth (*Choloepus hoffmanni*) (**Figure 1**).

CVe have previously been identified in the genome of the short-tailed opossum (*Monodelphis domestica*), an American marsupial [7]. In phylogenies based on rep, this sequence groups together with the porcine circoviruses, canine circovirus, and the CVe we identified in *Mus caroli*. We identified the first examples of CVe from the genomes of Australian marsupial species: the Tasmanian devil (*Sarcophilus harrisii*) and the koala (*Phascolarctos cinereus*). Both these sequences derived from circovirus *rep* genes, and grouped together in phylogenetic trees (**Figure S1**). However, their placement relative to other taxa was not supported with confidence, reflecting their short and degraded nature. Several other short and degraded matches to Rep probes were identified in other mammalian species (**Table 1, Table 2, Figure 1**). These sequences were relatively distantly related to one another and to contemporary circoviruses.

## 4. Discussion

### 4.1. CVe provide retrospective information about circovirus evolution

In this study, we recovered CVe from published vertebrate genomes, determined their genomic structures, and examined their phylogenetic relationships to contemporary circoviruses. Our analysis is the first to examine such a large set of CVe sequences, and to screen so widely within vertebrates. We show that CVe are relatively widespread in vertebrate genomes, though it appears they are absent from some lineages (e.g. primates, in which genome coverage is relatively high).

Several of the CVe loci identified here have been reported previously [7, 16, 19, 20], and the majority of novel CVe sequences recovered by our screen were orthologs or duplicates of these loci. Nevertheless, we identified 17 CVe loci that have not been reported before (**Table 1, Table S3**). These sequences provide the first evidence of (ancestral) circovirus infection for several species (**Table 2**). In addition, the identification and characterisation of novel orthologs allowed us to establish the first minimum age estimates for some CVe loci, and to markedly extended those of others. Thus, we were able to derive a more accurately calibrated timeline of evolution for the *Circovirus* genus, spanning multiple geological eras (**Figure 2**). Furthermore, we observed that CVe in fish, birds and mammals cluster phylogenetically with exogenous circoviruses identified from the same host class. This implies that there is a stability to the relationship between circovirus and host relationships, at least at higher taxonomic levels.

### 4.1. Impact of CVe on host genome evolution

The majority of CVe are derived from *rep* genes. To the extent that CVe have been exapted or co-opted, the predominance of CVe derived from *rep* might reflect that these sequences are more readily functionalised than those derived from *cap*. Furthermore, we identified one sequence. Notably one CVe in the mangrove rivulus (*Kryptolebias marmoratus*) encoded an intact *rep* gene that is predicted to express mRNA, suggesting it may have been functionalized in some manner (**Figure 1**).

Notably, several examples have now been described of endogenous viral elements (EVEs) that are derived from replicase genes, are expressed, and encode intact ORFs [7, 30, 31]. These elements are derived from a range of different viruses, and have clearly arisen in distinct events, suggesting there might be a common mechanism causing EVEs derived from the replicases of distinct viruses to be selected and maintained in different species. Alternatively, it is possible that the discrepancy in numbers simply reflects that *cap*-derived sequences are less conserved and therefore harder to detect.

Curiously, it is rare for more than one CVe to occur in the germline of any jawed vertebrate lineage. Carnivores are an obvious exception, since CVe have been amplified to relatively high copy number (10-20 copies) in several carnivore lineages (**Figure 4**), apparently via retrotransposon-mediated duplication. Further investigation of how these CVe have been amplified may reveal if their presence within an actively replicating retrotransposon lineage has impacted on the fixation of transposable elements derived from that lineage.

## Conclusions

We identified the complete repertoire of CVe sequences in published vertebrate genome assemblies. Through comparative analysis of these sequences, we provide the most complete picture yet of how viruses in the genus *Circovirus* have evolved and interacted with their hosts over the course of their evolution. The sequence-based resource implemented here can facilitate further characterisation of circovirus distribution, diversity and evolution as new CVe and circovirus sequence data become available.

## Acknowledgements

RJG was funded by the Medical Research Council of the United Kingdom (MC_UU_12014/12). WMS is supported by the Fundação de Amparo à Pesquisa do Estado de São Paulo, Brazil (Scholarships N0. 17/13981-0).

